# Blood flow-mediated gene transfer and siRNA-knockdown in the developing vasculature in a spatio-temporally controlled manner in chicken embryos

**DOI:** 10.1101/434068

**Authors:** Yuta Takase, Yoshiko Takahashi

**Author notes:** Author for correspondence: Prof. Yoshiko Takahashi, Department of Zoology, Graduate School of Science, Kyoto University.

## Abstract

We describe a method by which early developing vasculature can be gene-manipulated independently of the heart in a spatio-temporally controlled manner. Lipofectamine 2000 or 3000, an easy-to-use lipid reagent, has been found to yield a high efficiency of transfection when co-injected with *GFP* DNA within a critical range of lipid concentration. By exploiting developmentally changing patterns of vasculature and blood flow, we have succeed in controlling the site of transfection: injection with a lipid-DNA cocktail into the heart before or after the blood circulation starts results in a limited and widely spread patterns of transfection, respectively. Furthermore, a cocktail injection into the right dorsal aorta leads to transgenesis of the right half of embryonic vasculature. In addition, this method combined with the siRNA technique has allowed, for the first time, to knockdown the endogenous expression of *VE-cadherin* (also called *Cdh5*), which has been implicated in assembly of nasant blood vessels: when *Cah5* siRNA is injected into the right dorsal aorta, pronounced defects in the right half of vasculature are observed without heart defects. Whereas infusion-mediated gene transfection method has previously been reported using lipid reagents that were elaborately prepared on their own, Lipofectamine is an easy-use reagent with no requirement of special expertise. The methods reported here would overcome shortcomings of conventional vascular-transgenic animals, such as mice and zebrafish, in which pan-endothelial enhancer-driven transgenesis often leads to the heart malformation, which, in turn, indirectly affects peripheral vasculature due to flow defects. Since a variety of subtypes in vasculature have increasingly been appreciated, the spatio-temporally controllable gene manipulation described in this study offers a powerful tool.

**Research Highlights:**

- Blood flow-mediated transfection enables site-specific transgenesis in vessels.
- This transfection technique allows local knockdown of endogenous gene(s) by siRNA.
- Knockdown of endogenous *VE-cadherin* causes vascular defects without heart failure.

## Introduction

Blood vessel networks are distributed widely in the vertebrate body, and they deliver essential substances including oxygen, nutrients, and hormones to tissues and organs (Adams and Alitalo, 2007; Potente et al., 2011). The vascular system also plays important roles in a variety of functional events such as homeostasis, tissue morphogenesis, regeneration, and cancer metastasis (Carmeliet, 2005; Makita et al., 2008; Saito et al., 2012). During blood vessel formation in embryogenesis, endothelial precursor cells called angioblasts emerge in mesenchymal populations, and subsequently differentiate into endothelial cells to form nascent blood vessels by the process of vasculogenesis (Herbert and Stainier, 2011). Following vasculogenesis, blood vessels undergo angiogenesis, during which time they change their structures by extending, sprouting, pruning, and remodeling (Adams and Alitalo, 2007; Herbert and Stainier, 2011). Finally, mature vasculatures become associated with type-specific perivascular cells (Jain, 2003) to complete arterial-venous differentiation.

An increasing body of experimental evidence has advanced our understanding of the molecular and cellular mechanisms by which blood vessel formation is regulated. The major approaches used for the analyses have been the endothelial-specific loss- or gain-of-function of genes, for which transgenic animals in mice and zebrafish serve as powerful experimental models (De Val and Black, 2009; Hogan and Schulte-Merker, 2017; Stahl et al., 2010). In these experiments, pan-endothelial drivers, such as Tie1 or Tie2, are often used, and this leads to a transgenesis of the entire cardiovasculature system, including both the heart and vessels (Korhonen et al., 1995; Schlaeger et al., 1997). If heart development is affected by the transgene (for example by interfering with ALK5 or Smad4 expression), then blood flow is frequently undermined; and this loss of mechanical flow can indirectly cause the malformation of peripheral vasculature (De Val and Black, 2009; Lan et al., 2007; Pardanaud et al., 1996; Sridurongrit et al., 2008). This makes it difficult to distinguish between the direct and indirect effects of transgenesis in peripheral vasculatures.

Recent reports have shown that the vasculature contains numerous subtypes of blood vessels. There is not only the artero-venus specification, but also the region-specific diversity of shapes and functions of vessel branches, which are being increasingly appreciated (Augustin and Koh, 2017; Pardanaud et al., 1996; Sato et al., 2008). However, how these diverse types of peripheral vasculature are established remains poorly explored, mainly because methods that allow region-specific gene manipulations of vasculature have been limited.

Early chicken embryos are flat in structure, in which a variety of vascular subtypes can be seen, including dorsal aortae, intersegmental vessels, and yolk sac vasculature as early as at embryonic day 2 (E2). Rosselo and Torres (2010) reported that the *in ovo* electroporation into developing chick dorsal aortae resulted in local transgenesis (Rossello and Torres, 2010). However, as we confirmed (unpublished data), the efficiency of such gene transfer is low. Other groups (Bollerot et al., 2006; Decastro et al., 2006) have reported techniques of liposome-mediated transfection into chicken embryonic vasculature, but these methods require high expertise of liposome construction and preparation. So far, no method has been reported to knockdown endogenous gene(s) in chicken vasculature.

Here, we demonstrate methods that enable efficient and local transgenesis of vasculature in chicken embryos using the commercially available reagent Lipofectamine, which has been widely and commonly used for gene transfection into cultured cells. Following infusion with a cocktail containing Lipofectamine and *GFP* cDNA, GFP-positive endothelial cells were found in a variety of vasculature subtypes. By exploiting blood flow and vascular patterns that change in a stage-dependent manner, we were able to control transfected areas by changing the site and time point of injection along the developmental stages. In addition, our transfection technique allowed local knockdown of the endogenously expressed *VE-cadherin* (also called *Cdh5*) gene by small interfering RNA (siRNA). The methods described here allow us to spatially and temporally regulate gain-of-function and loss-of-function gene expression in the developing vasculature, without requiring complicated liposome preparation protocols or transgenic animals.

## Materials and Methods

### Embryological manipulations

Fertilized chicken eggs were commercially obtained from the poultry farm Shiroyama Farm (Kanagawa, Japan). Embryos were staged according to Hamburger & Hamilton (1951) or somite stage (ss). All animal experiments were conducted with the ethical approval of Kyoto University.

### Expression Vectors

pCAGGS-GFP and pCAGGS-mCherry plasmids were described previously (Momose et al., 1999; Yokota et al., 2011). The open reading frame (ORF) of chicken *Cdh5* was PCR-amplified using following primers:

Cdh5-Fw, 5’-CCACCGGTCGCCACCATGAAGAAGCTTATTCTGCT-3’;
Cdh5-Rv, 5’-GCCCTTGCTCACCATCTCGAGGGAATACACAAAATCTTCAT-3’.

pCMV-Cdh5GFP plasmid was made by fusing the Cdh5 ORF to pCMV-GFP (Clontech) using In-Fusion HD Cloning Kits (Clontech). pCMV-GFP linearized vector was amplified from pEGFP-N1 (Clontech) using following primers:

GFP-N1-Fw, 5’-ATGGTGAGCAAGGGCGAGGA-3’;
GFP-N1-Rv, 5’-GGTGGCGACCGGTGGATCCC-3’.

Cdh5mutGFP, which was resistant to Cdh5-siRNA#1, was made using PrimeSTAR Mutagenesis Basal Kit (Takara) with following primers (capital letters indicate replaced nucleotides):

Cdh5mut-Fw, 5’-atcTTTAgaGccTccTAGCaaGtttattatcaaggtttctgat-3’
Cdh5mut-Rv, 5’-aaaCttGCTAggAggCtcTAAAgatcggttgtttcttctgtca-3’

### siRNAs

We designed siRNAs to interfere with *Cdh5* expression, referring to technical information of BLOCK-iT RNAi Designer (Invitrogen) and siRNA Target Finder (Genscript). The following 19-mer sequences were selected:

Cdh5-siRNA#1 (at positions 369-387 of ORF), 5’-GCTGGAACCACCATCTAAA-3’;
Cdh5-siRNA#2 (at positions 1052-1070 of ORF), 5’-CAACAATTACCATTGAAGT-3’;
Cdh5-siRNA#3 (at positions 1470-1488 of ORF), 5’-GGTAATCATCAGGATTTCA-3’;
Luc-siRNA (negative control interfering with Luciferase expression), 5’-GGATCCTATCCGAAGGCAA-3’.

All sense and anti-sense RNA oligos were synthesized by Eurofins Genomics. Annealing of sense and anti-sense RNA oligos was prepared according to the manufacturer’s instruction. Rhodamine labeling of siRNA (Cdh5-siRNA#1) were made using *Label* IT siRNA Tracker Kit (TAKARA).

### mRNAs

*Luciferase* and *Cdh5mut* capped mRNAs were prepared from pBS-Luciferase and pBS-Cdh5mut, respectively, using T7 mScript Standard mRNA Production System (CellScript).

### Cell culture

Chicken fibroblast-derived DF1 cells were maintained with Dulbecco’s Modified Eagle Medium (DMEM) supplemented with 10% (v/v) fetal bovine serum (FBS) at 38.5°C, 5% CO_2_. For evaluation of Cdh5-siRNA efficacy, pCMV-Cdh5GFP (1,800 ng) and pCAGGS-mCherry (1,000 ng) were co-transfected into DF1 cells along with 30 pmol of each siRNA using Lipofectamine 2000 (Invitrogen). Twenty-four hours after transfection, DF1 cells were fixed overnight in phosphate buffered saline (PBS) containing 4% paraformaldehyde (PFA) at 4°C. Fluorescent images were obtained using AZ-C1 macro-confocal microscope system (Nikon). For cell counting analyses, images were processed using ImageJ (NIH).

### Visualization of blood vessel

*In vivo* visualization of blood vessels was performed by infusion with fluorescent ink as previously described (Takase et al., 2013). Chicken embryos were infused with 0.5-1 μl highlighter ink (PILOT spotliter; 1:10 dilution in PBS) through the heart using a micropipette pulled from a glass capillary (Narishige, GD-1) with a vertical micropipette puller (Narishige, PC-10). Manipulated embryos were incubated at 38.5°C for an additional period of 5 min before harvest. Fluorescent images were obtained using the Leica MZ10 F microscope (Leica) with the DS-Ri1 camera (Nikon).

### *In vivo* transfection into embryonic vasculature

For plasmid DNA transfection, Lipofectamine 2000 (Invitrogen), Lipofectamine 3000 (Invitrogen), ViaFect (Promega), FuGENE HD (Promega) and SuperFect (QIAGEN) were used. Each transfection reagent was diluted with Opti-MEM 25, 50, 75 or 100% (v/v), and plasmid DNA solution in Opti-MEM in was separately prepared at 600 ng/μl. Subsequently, the diluted transfection reagent and solution were mixed at a final concentration of 0, 12.5, 25, 37.5 or 50% (v/v) and 300 ng/μl, respectively, and incubated at room temperature for 5 min to allow the lipid– DNA complex to form.

Using a glass micropipette, 0.8 μl of the lipid–DNA complex solution was injected into the heart or the right dorsal aorta (R-DA) at the appropriate embryonic stages. For siRNA transfection, Lipofectamine 2000, siRNA and BLOCK-iT Alexa Fluor Red Fluorescent Control (Invitrogen) were diluted with Opti-MEM at a final concentration of 12.5% (v/v), 5 μM and 2.5 μM, respectively. For siRNA and mRNA transfection, Lipofectamine 2000, rhodamine-labeled Cdh5-siRNA#1 and *Luciferase* or *Cdh5mut* mRNA were diluted with Opti-MEM at a final concentration of 12.5% (v/v), 4 μM and 360 ng/μl (*Luciferase* mRNA) or 600 ng/μl (*Cdh5mut* mRNA), respectively.

### RNA probe and whole-mount *in situ* hybridization

cDNA fragment of chicken *Cdh5* was as described (Sato et al., 2008). Digoxigenin-labeled RNA probe was prepared according to the manufacturer’s instruction (Roche). Whole-mount *in situ* hybridization was performed as previously described (Tonegawa et al., 1997).

### Frozen sections

Chicken embryos were fixed overnight in PBS containing 4% PFA at 4°C. Frozen sections (10 μm thick) of fixed embryos were prepared with a cryostat (MICROM, HM500 OM). The sections were washed in PBS three times (each 5 min), and were sealed by FluorSave reagent (Calbiochem) containing 4′6′-diamidino-2-phenylindole dihydrochloride (DAPI). Fluorescent images were obtained using Axioplan 2 microscope with Apotome system (Carl Zeiss).

## Results

### Reagents for *in vitro* transfection can be used for vascular transgenesis in live, intact chicken embryos

Lipofectamine 2000 (Invitrogen) is a widely used reagent for the *in vitro* transfection into cultured cells. We asked if this reagent could also be used for *in vivo* transfection to target developing vasculatures in chicken embryos. We injected 0.8 μl of transfection cocktail (a solution containing Lipofectamine 2000 (lipid) and DNA) into the heart at embryonic day 2 (E2; HH14) at 5 different concentrations of lipid, 0%, 12.5%, 25%, 37.5%, and 50%, along with 300 ng/μl of pCAGGS-GFP (Fig. 1A). After 17 hours, relatively high levels of GFP signals were observed in the heart and nearby vessels in embryos that received the lipid concentration of 12.5% or 25% (Fig. 1C, n = 25; Fig. 1D, n = 40, Table 1), whereas a cocktail at concentrations of 37.5% or 50% resulted in barely detectable/no GFP expression or embryonic death (Fig. 1E, F, n = 10 each, Table 1). Expression levels of GFP were categorized into three classes: no expression (-), moderate (+), and high (++). The highest GFP level with the lowest frequency of embryonic death was achieved with the concentrations of 12.5% or 25% of Lipofectamine 2000 (Table 1).

**Figure 1.**
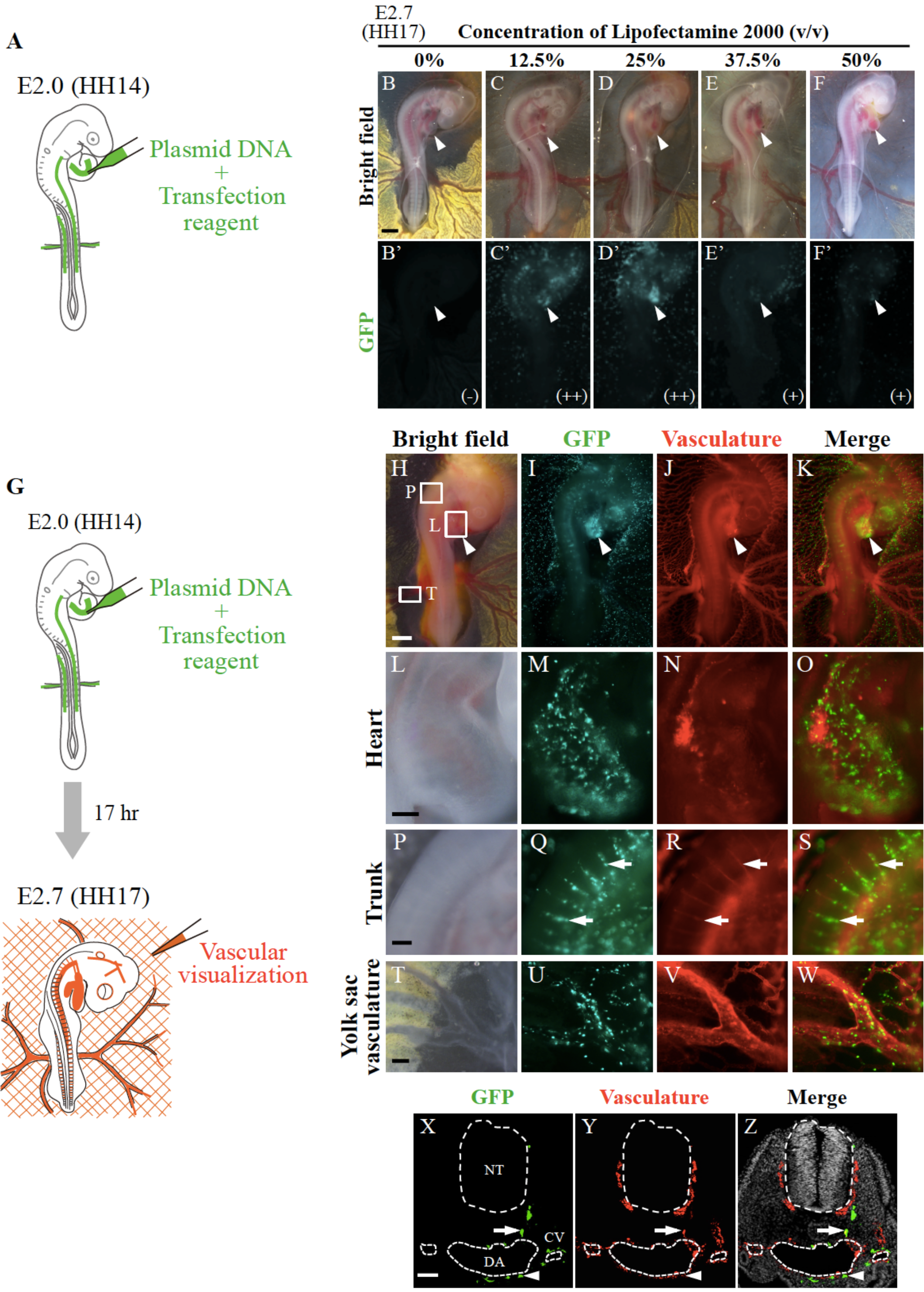
Commercially available reagents can be used for vascular transgenesis in chicken embryos. (A) A diagram showing injection with a liposome-DNA cocktail. A cocktail containing pCAGGS-GFP plasmid (300 ng/μl) and transfection reagent, shown in green, was injected into the heart of E2 (HH14) embryos, and GFP signals were assessed 17 hour after transfection (E2.7; HH17). Transfection efficiency was classified into three levels (-, +, ++) based on GFP expression in the heart. (B–F) GFP expression in transfected embryos at E2.7. Embryos receiving the concentration of Lipofectamine 2000 at 12.5% and 25% exhibited high level of GFP expression (++), whereas embryos receiving 37.5% and 50% of Lipofectamine displayed a moderate level (+). (G) A diagram showing injection with a liposome-DNA cocktail and vascular visualization. A cocktail containing pCAGGS-GFP plasmid (300 ng/μl) and Lipofectamine 2000 (12.5%), shown in green, was injected into the heart of E2 (HH14) embryos. Vascular networks of transfected embryos were visualized by perfusion with fluorescent ink at E2.7 (HH17) shown in red. (H–W) GFP expression and vascular networks of transfected E2.7 embryo. (H-K) Widespread GFP expression was observed in the vasculature, and a high level of GFP signals was detected in the heart (arrowheads), into which the lipid–DNA cocktail was injected. (L-W) GFP signals in the heart (L-O), intersegmental vessels (arrows) (P-S), and the yolk sac vasculature (T-W). (X-Z) Transverse sections of the trunk region of transfected embryo at E2.7. GFP signals were detected in endothelial cells labeled with infused violet ink. The arrowhead and arrow indicate endothelial cells of the DA and ISV, respectively. NT, neural tube; DA, dorsal aorta; CV, cardinal vein. Scale bars: 1 mm for (B-F, H-K), 250 μm for (L–W), and 50 μm for (X-Z).

**Table.**
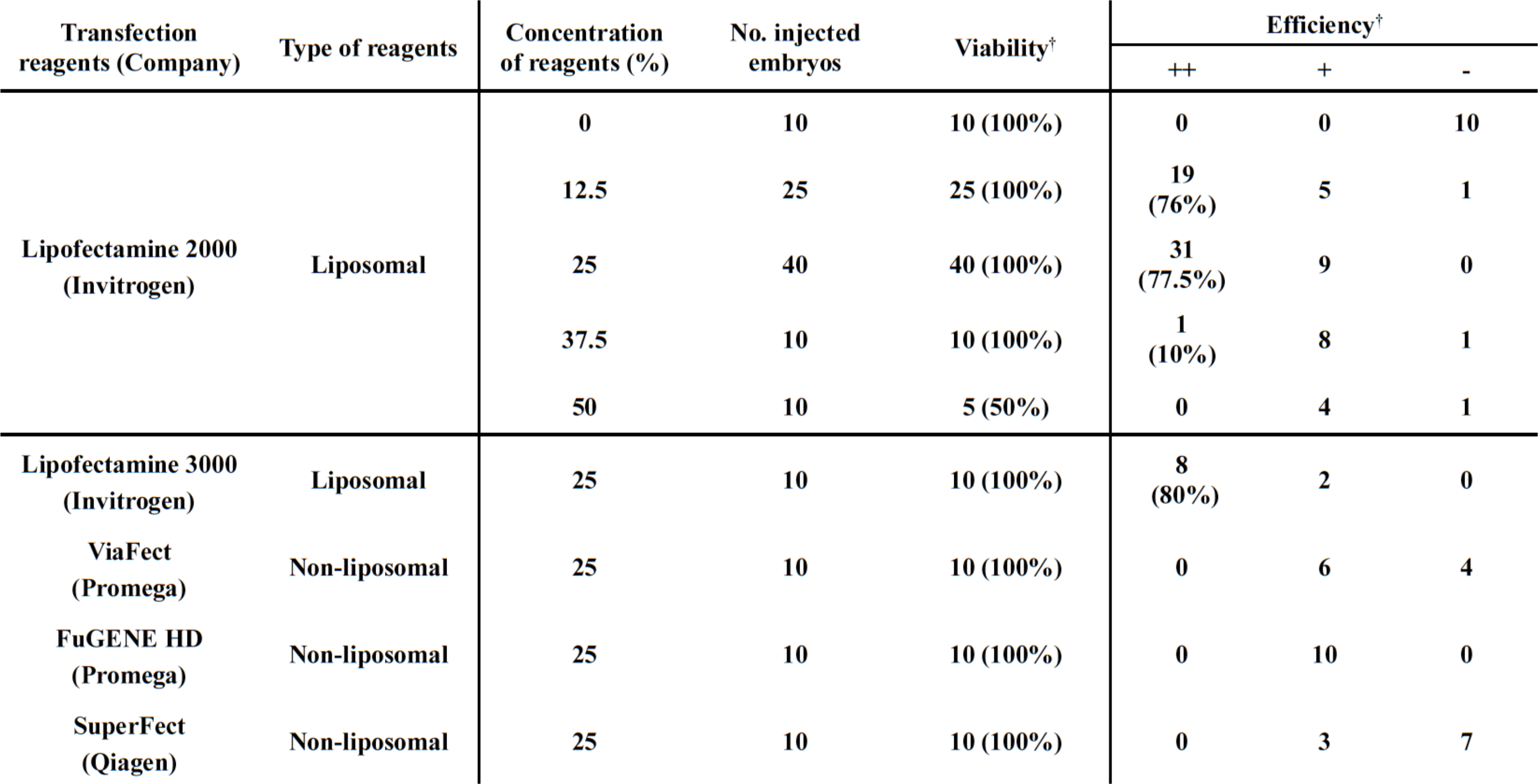
Comparison of transfection efficiencies among commercial transfection reagents. A GFP-cocktail with each of transfection reagents (Lipofectamine 2000, Lipofectamine 3000, ViaFect, FuGENE HD, and SuperFect) was injected into the heart at HH14, and GFP signals were assessed after 17 hrs in the heart. See text for details. -, no GFP expression; +, low or moderate GFP expression; ++, high GFP expression.

We also tested other commercially available reagents for transfection, Lipofectamine 3000 (Invitrogen), ViaFect (Promega), FuGENE HD (Promega), and SuperFect (Qiagen) at the concentration of 25%. Among these, only Lipofectamine 3000 yielded GFP signals comparable to those by Lipofectamine 2000 (Table 1). For further studies, we used Lipofectamine 2000 at 12.5%, since it is less expensive than Lipofectamine 3000.

To further scrutinize transfected sites in embryos, we visualized blood vessels of cocktail-injected embryos by ink perfusion prior to harvest 17 hours after transfection (Fig. 1G) (Takase et al., 2013, see also Materials and Methods). In addition to the heart, GFP signals were widely distributed in blood vessels including intersegmental vessels (ISVs) and yolk sac vasculature (YS) (Fig. 1H-W). Circulating blood cells were also transfected (data not shown). The endothelial transfection was also confirmed in transverse sections: Fig. 1X-Z show GFP-positive endothelial cells in the dorsal aorta (DA), ISV, and cardinal vein. Thus, Lipofectamine 2000 can be used for vascular transgenesis in chicken embryos.

### Site-specific transgenesis of developing vasculature

We examined whether GFP-transfected sites in the vasculature would differ if the developmental stage and site of cocktail injection were changed. When the lipid-DNA cocktail is injected into a beating heart, the cocktail solution must be delivered by blood flow. In contrast, if injected into an early-formed heart before the circulation starts, the cocktail would spread only locally. Likewise, the delivery pattern of the cocktail must be different if injected into one of paired dorsal aortae. In this way, we compared GFP-transfected areas in vasculature by testing three different developmental stages, 13 ss, 16 ss and 19 ss, and two different sites of injection, the heart and right dorsal aorta (R-DA) (Fig. 2). Ink was perfused prior to harvest to confirm that the transfected cells were confined to endothelial cells.

**Figure 2.**
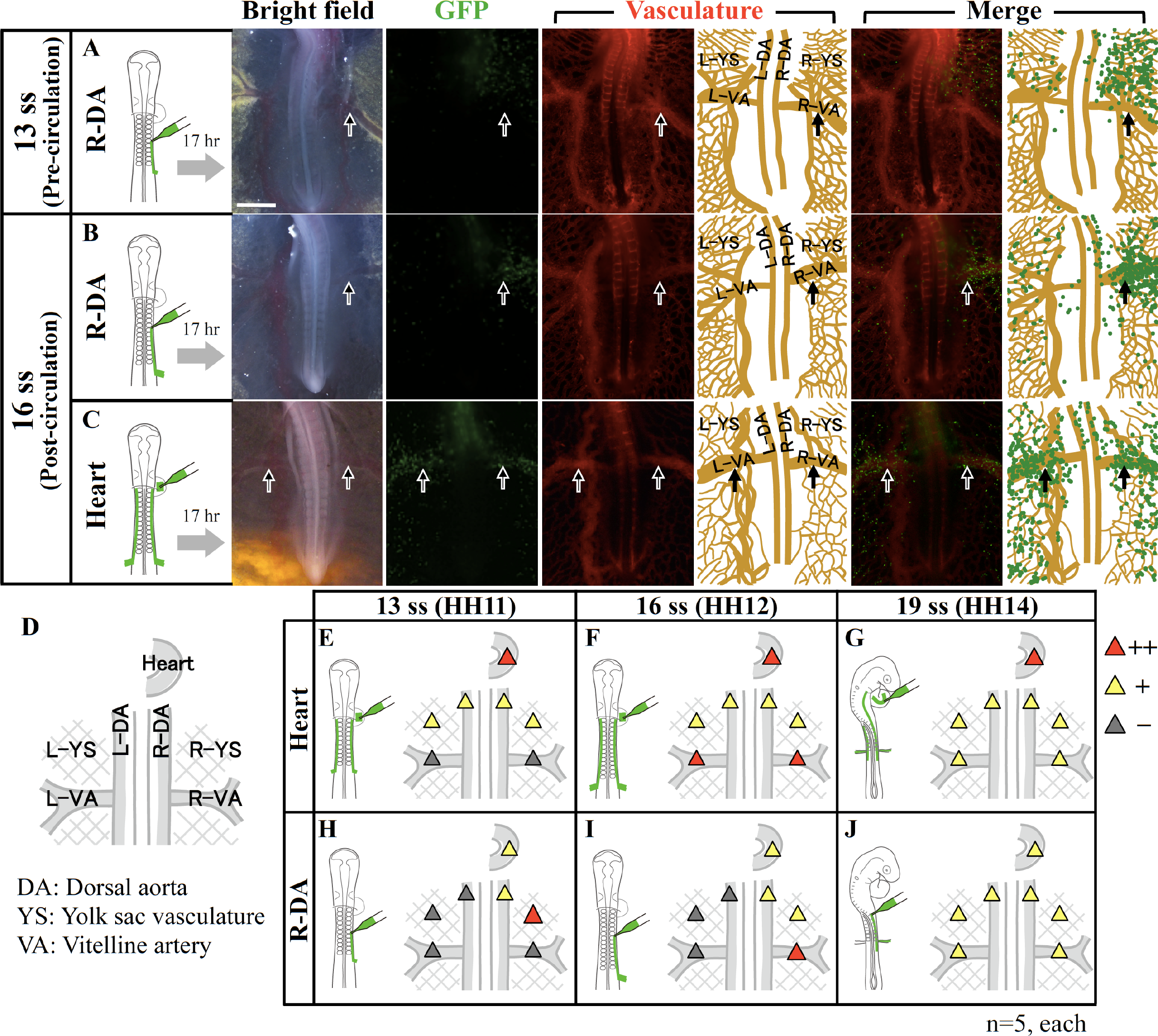
Site-specific transgenesis of developing vasculature by adjusting developmental stages and injection sites. (A–C) A cocktail of pCAGGS-GFP plasmid and Lipofectamine 2000 was injected into the right dorsal aorta (R-DA) or heart at two different stages, 13 ss and 16 ss. GFP-signals were assessed 17 hour after transfection. Black arrows indicate vitelline arteries. (A) When a cocktail was injected into R-DA at 13 ss of pre-circulation stage), GFP-transfected area was in a limited range within the anterior YS of the right side of embryo (R-YS) and R-DA. (B) When injected into R-DA at a more advanced stage of 16 ss, the GFP-transfected area was expanded with R-DA, R-YS and R-VA being labeled. (C) When injected into the heart of 16 ss embryo, the GFP-positive area was seen in both sides of embryo including DAs, YSs, and VAs. Scale bar: 1 mm. (D-J) Summary of GFP-transfection in 7 different areas (D) after cocktail injections in two different positions (heart and R-DA) and three stages (13 ss, 16 ss, 19 ss). GFP signal intensities were classified into three levels: high expression (red, ++), moderate expression (yellow, +), and low or no expression (grey, -).

When injected into the R-DA at the pre-circulation stage (13 ss; no blood flow), an GFP-positive area was within a limited range in the anterior YS of the right side of embryo (R-YS) (Fig. 2A, H). In contrast, when injected into the R-DA after the circulation started (16 ss), the GFP-positive area was expanded, and the right vitelline artery (R-VA) was highly labeled (Fig. 2B, I). When injected into the heart at the same stage (16 ss), the transfected area was seen in both sides of embryo including YSs and VAs (Fig. 2C, F).

Fig. 2D-J summarize relative levels of GFP expression assessed in 7 different areas of embryos, the heart, right and left DAs, right and left YSs, and right and left VAs, when the cocktail was injected into different stages and different sites; three different embryonic stages of 13 ss (pre-circulation), 16 ss (onset of circulation), and 19 ss (circulation established), and two different injection sites, the heart and R-DA, were tested.

When injected into the heart of pre-circulation (13 ss), the cocktail remained in limited areas of YSs of both sides of embryos (Fig. 2E). In contrast, when injected into the heart after the circulation started (16 ss), the cocktail spread widely, and GFP expression was preferentially observed in VAs, in which blood flow was predominant compared to their neighboring vascular plexus (capillaries) (Fig. 2F). After the establishment of blood circulation, the cocktail injected into the heart spread over the entire vasculature (Fig. 2G, GFP-positive cells in DA, YS, VA are shown). When injected into the right DA, the right half of vasculature exhibited sequential changes in GFP distribution which was similar to those in the case of the heart injection (Fig. H-J). Together, site-specific transfection in early chicken vasculature is enabled by adjusting both developmental stages (pre-circulation vs post-circulation) and injection sites of a lipid–DNA cocktail.

### siRNA-mediated *Cdh5* knockdown: *In vitro* assay for evaluation of efficiency

To test if the aforementioned method could readily accomplish siRNA-mediated gene knockdown in chicken embryonic vasculature, we used the *VE-Cadherin* gene (also called *cadherin 5* (*Cdh5*), which is implicated to be important for the assembly of nascent blood vessels in mice (Crosby et al., 2005). We prepared three siRNAs against the *Cdh5* ORF (Cdh5-siRNA#1, #2, #3) (Fig. 3A).

**Figure 3.**
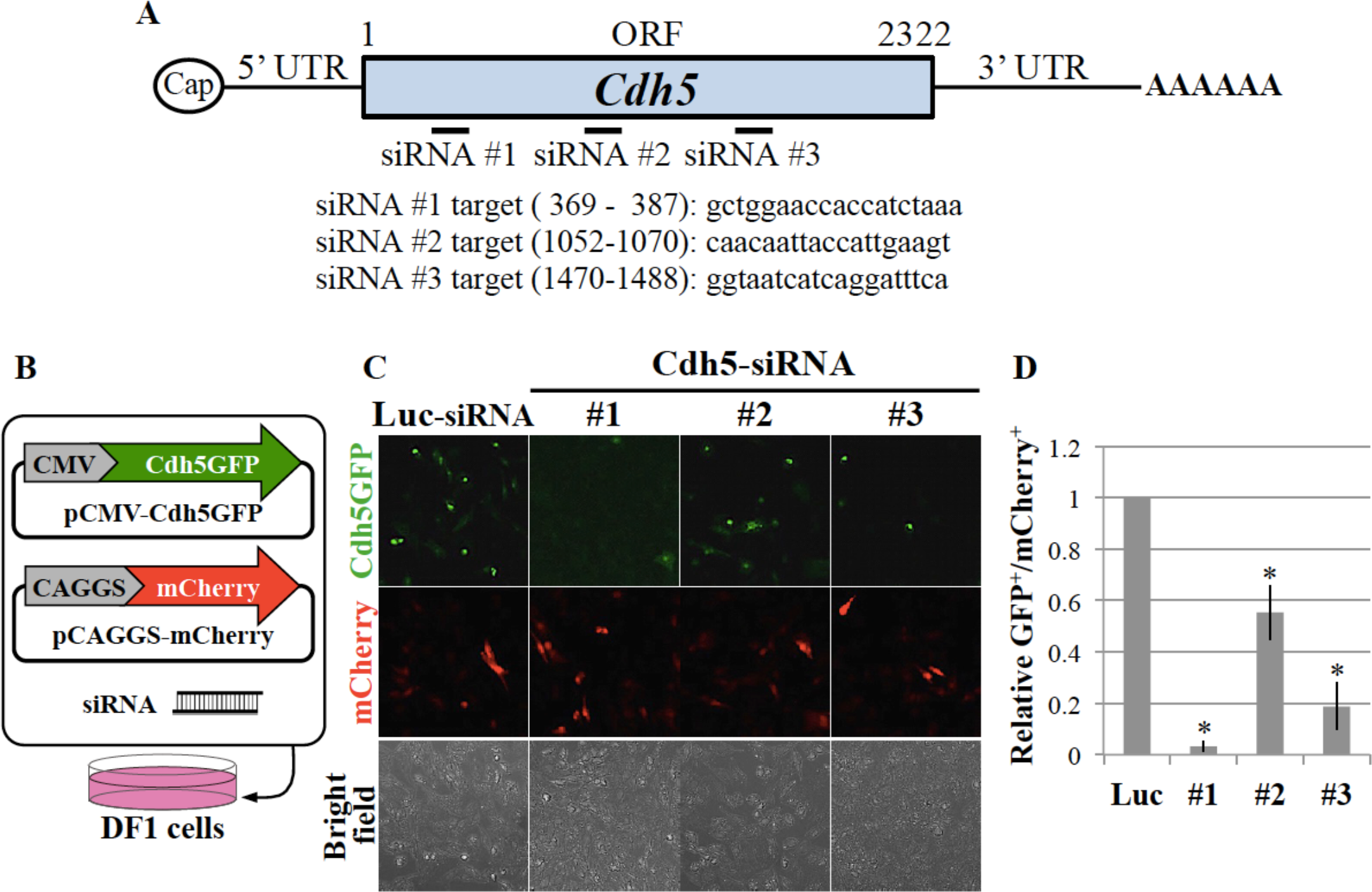
*In vitro* evaluation of siRNAs against *Cdh5*. (A) Schematic structure of chicken *Cdh5* mRNA and siRNA-targeted regions. Three siRNAs against *Cdh5* (Cdh5-siRNA#1, #2 and #3) were designed to target the open reading frame (ORF). (B) pCMV-Cdh5GFP and pCAGGS-mCherry plasmids were co-transfected into DF1 cells with each of Luc-siRNA or Cdh5-siRNAs#1–3, and GFP intensity was assessed 24 hrs after transfection. (C) In Cdh5-siRNA#1–3 co-transfected DF1 cells, whereas the number of mCherry^+^ cells was comparable to that in Luc-siRNA co-transfected cells, the number of Cdh5GFP^+^ cells markedly decreased receiving Cdh5-siRNA#1 or -#3. (D) Quantitative representation of a relative ratio of the number of Cdh5GFP^+^ cells to the number of mCherry^+^ cells (n = 4 each). Error bars represent SD (standard deviation). \**P* < 0.001.

To evaluate the knockdown efficiency by these Cdh5-siRNAs, we performed *in vitro* assay using chicken fibroblast-derived DF1 cells. The cells were co-transfected with expression vectors of GFP-tagged Cdh5 (Cdh5GFP) and mCherry, the latter being used as an internal control of transfection efficiency, along with each of three Cdh5-siRNAs (Fig. 3B). Luciferase-siRNA (Luc-siRNA) was used as a negative control. DF1 cells co-transfected with Cdh5GFP, mCherry, and Luc-siRNA displayed both GFP and mCherry signals (Fig. 3C). In contrast, intensity of GFP signals was reduced when co-transfected with Cdh5-siRNA: a relative ratio of the number of Cdh5GFP^+^ cells to that of mCherry^+^ cells (normalized by the index of Luc-siRNA control) was 0.03 ± 0.02 siRNA#1, 0.55 ± 0.11 for Cdh5-siRNA#2, and 0.19 ± 0.09 for Cdh5-siRNA#3 (Fig. 3C, D, n=4 for each). These results indicate that Cdh5-siRNAs prepared in this study could inhibit expression of exogenously introduced Cdh5GFP in culture cells, with Cdh5-siRNA#1 exerting the highest efficiency.

### Site-specific gene knockdown in developing vasculature

We next asked whether our flow-mediated gene transfer method was applicable for knocking down endogenous *Cdh5*, which would also be expected to cause vascular defects. The three Cdh5-siRNAs were separately injected into the right dorsal aorta (R-DA) of 16 ss (HH12) embryos as a 0.8 μl lipid cocktail (12.5% of Lipofectamine 2000, 5 μM of each siRNA, and 2.5 μM of Alexa Fluor Red Fluorescent Control, a fluorescent tracer of siRNA transfection) (Fig. 4A). When the embryos developed to HH16 (20 hrs post injection), vascular formation was assessed by yellow ink infusion. siRNA-transfected sites were confirmed by co-injected Alexa Fluor Red Fluorescent signals.

**Figure 4.**
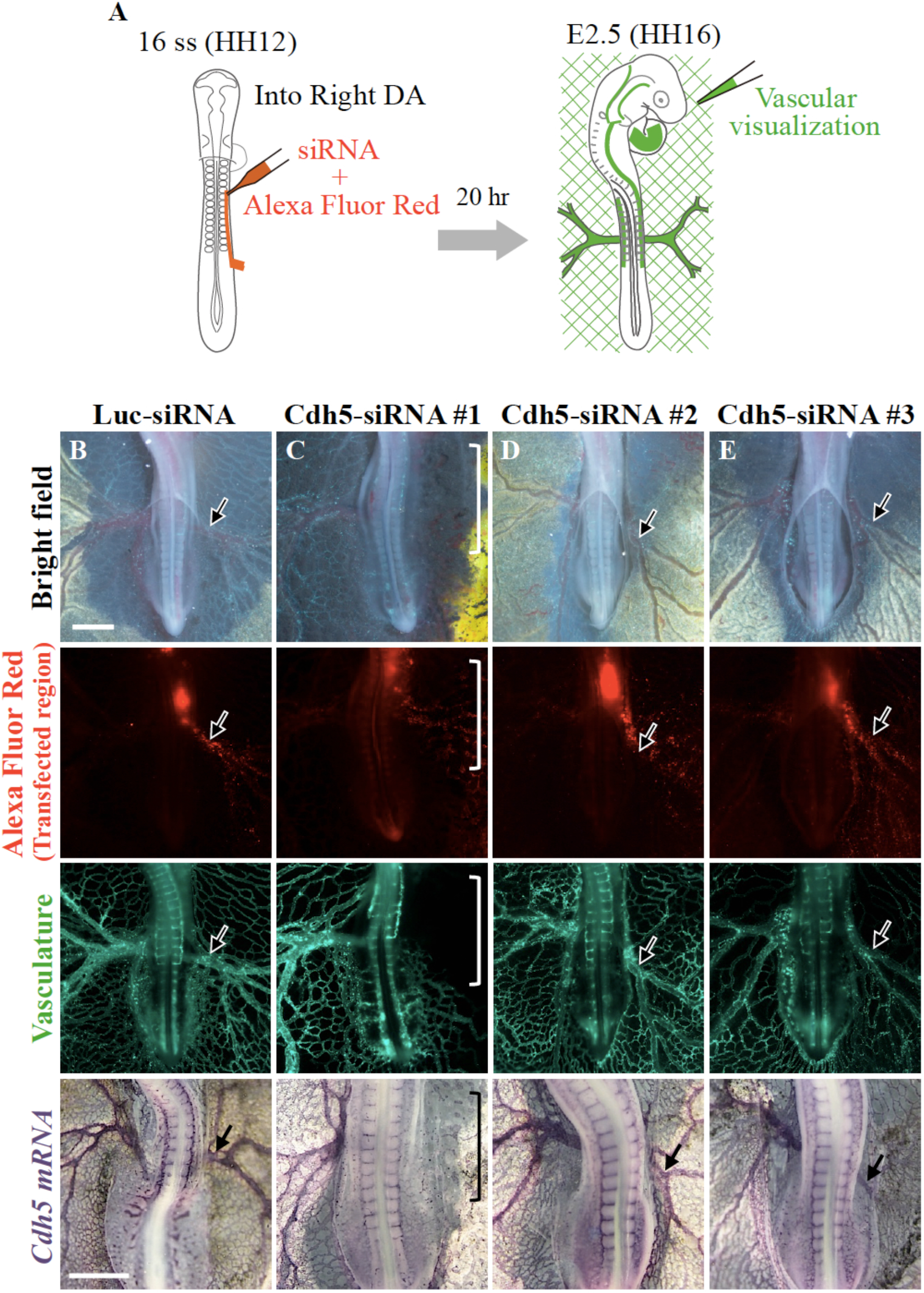
Site-specific *Cdh5* knockdown in developing vasculature caused local disruption of vascular network. (A) A cocktail containing siRNA, Alexa Fluor Red, and Lipofectamine 2000 was injected into the R-DA of 16 ss embryos, followed by examination 20 hrs after transfection (E2.5). Vascular networks of E2.5 (HH16) embryos were visualized by perfusion with fluorescent ink, shown in green. (B–E) Phenotypes of embryos transfected with siRNAs (shown on the top). Bright field, Alexa Fluor Red, and vasculature photos were taken of the same embryos. Specimens for *in situ* hybridization with *Cdh5* mRNA were different from those shown above. Arrows indicate R-VA. (B) Luc-siRNA-transfected embryos showed normal vascular networks (n=15/16). (C) Cdh5-siRNA#1-transfected embryos exhibited a local disruption of vasculature corresponding to the transfected area in the right side including R-YS and R-VA (brackets) (n=29/32). (D, E) Embryos transfected with Cdh5-siRNA#2- or -#3 displayed mild (#3) or almost undetectable (#2) phenotypes compared to those with Cdh5-siRNA#1 (n=17 each). R-DA, right dorsal aorta; R-VA, right vitelline artery. Scale bar: 1 mm.

We found that the formation of R-YS and R-VA was markedly affected, as revealed by both ink perfusion and *in situ* hybridization with *Cdh5* mRNA (Fig. 4C, n = 29/32), whereas effects on the vasculature in the left side (L-YS and L-VA) were unrecognizable. Consistent with differential knockdown efficiencies among the three Cdh5-siRNAs (Fig. 3), embryos injected with Cdh5-siRNA#1 displayed most pronounced defects in vasculature, with siRNA#2 and siRNA#3 showing little and moderate activities, respectively (Fig. 4C-E; #1: n=32, #2 and #3: n=17). Vasculatures injected with Luc-siRNA-containing cocktail were indistinguishable from those of untreated embryos (Fig. 4B, n = 15/16).

Together, these studies provide proof-of-concept that the blood flow-mediated transfection *in vivo* with Lipofectamine 2000 can be used for siRNA-knockdown. Importantly, this method allows a site-specific knockdown in vasculature, which has been difficult by conventional genetics such as using Tie2-Cre/flox mice.

When the Cdh5-siRNA#1 cocktail was injected into a beating heart, the entire vasculature, including the heart, was severely affected, resulting in malformed embryos (n=18 for Luc-siRNA; n=16 for Cdh5-siRNA#1; Fig. 5). Such embryo-wide malformation is reminiscent of phenotypes reported for *Cdh5*-knockout mice, which die at mid-gestation as a result of severe cardiovascular anomalies, including both vascular and cardiac defects (Carmeliet et al., 1999; Gory-Faure et al., 1999) making it difficult to distinguish between direct and indirect effects by the gene knockdown. Thus, the flow-mediated site-specific knockdown described in the current study offers significant merits.

**Figure 5.**
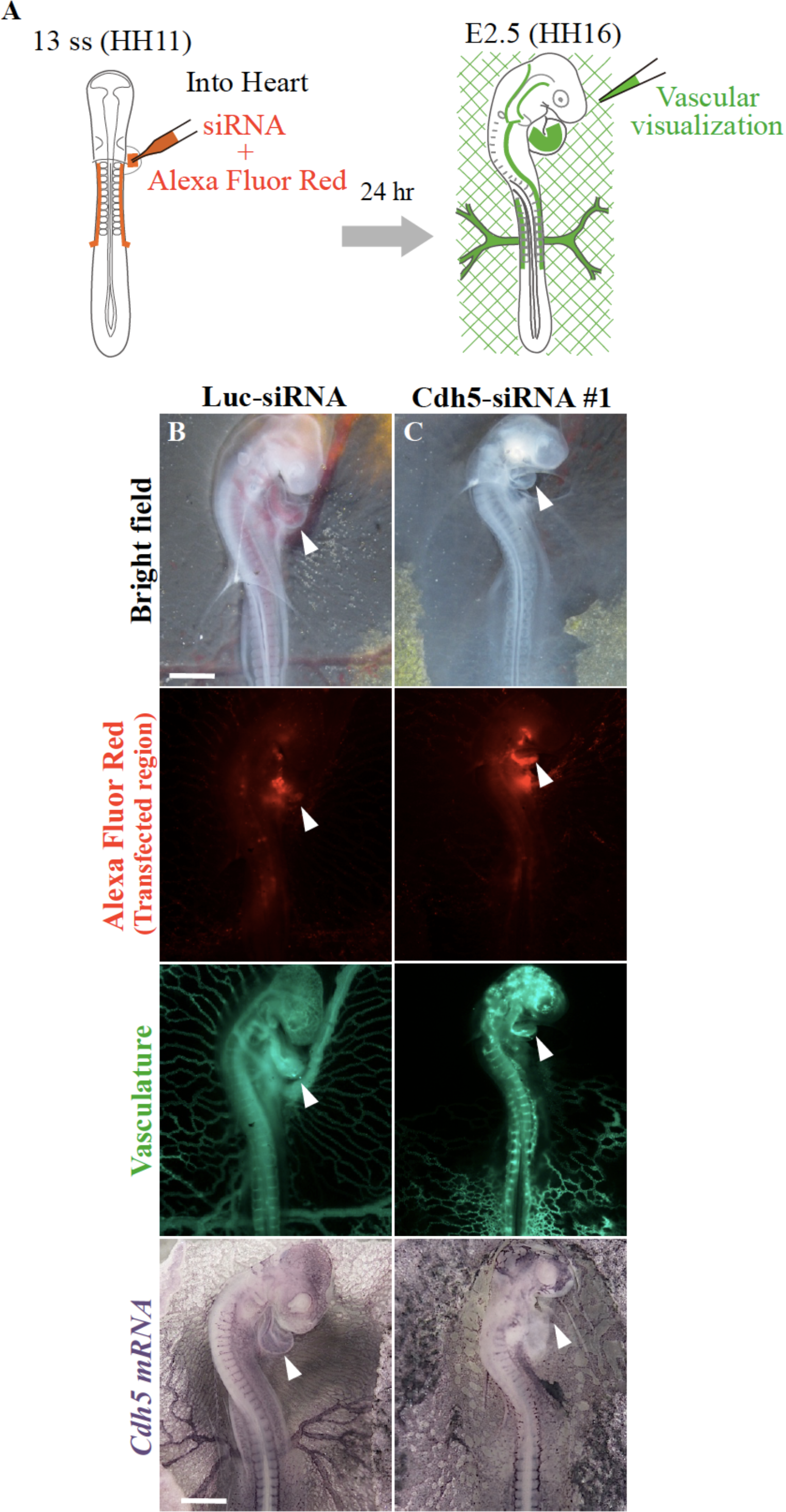
Cdh5 knockdown in the heart caused heart defects (arrowheads) and embryonic malformation with severe defects in vasculature. (A) A cocktail containing siRNA, Alexa Fluor Red, and Lipofectamine 2000 was injected into the hearts of 13 ss embryos, which were examined after 24 hrs at E2.5 with ink perfusion (green). (B, C) Bright field, Alexa Fluor Red, and vasculature photos were taken of the same embryos. Specimens for *in situ* hybridization with *Cdh5* mRNA were different from those shown above. Whereas Luc-si-transfected embryos were normal (n=15/18), embryos transfected with Cdh5-siRNA#1 exhibited were malformed with severe vascular defects (n=12/16). *Cdh5* mRNA was only minimally detected in the heart of Cdh5-siRNA#1 transfected embryos. Scale bar: 1 mm.

### Rescue of Cdh5 siRNA-knockdown by co-transfection with *Cdh5* mRNA

To determine if the vascular defects observed in the siRNA#1-injected embryos were elicited specifically by Cdh5-specific knockdown, we performed rescue experiments using siRNA-resistant *Cdh5* mRNA (Fig. 6).

**Figure 6.**
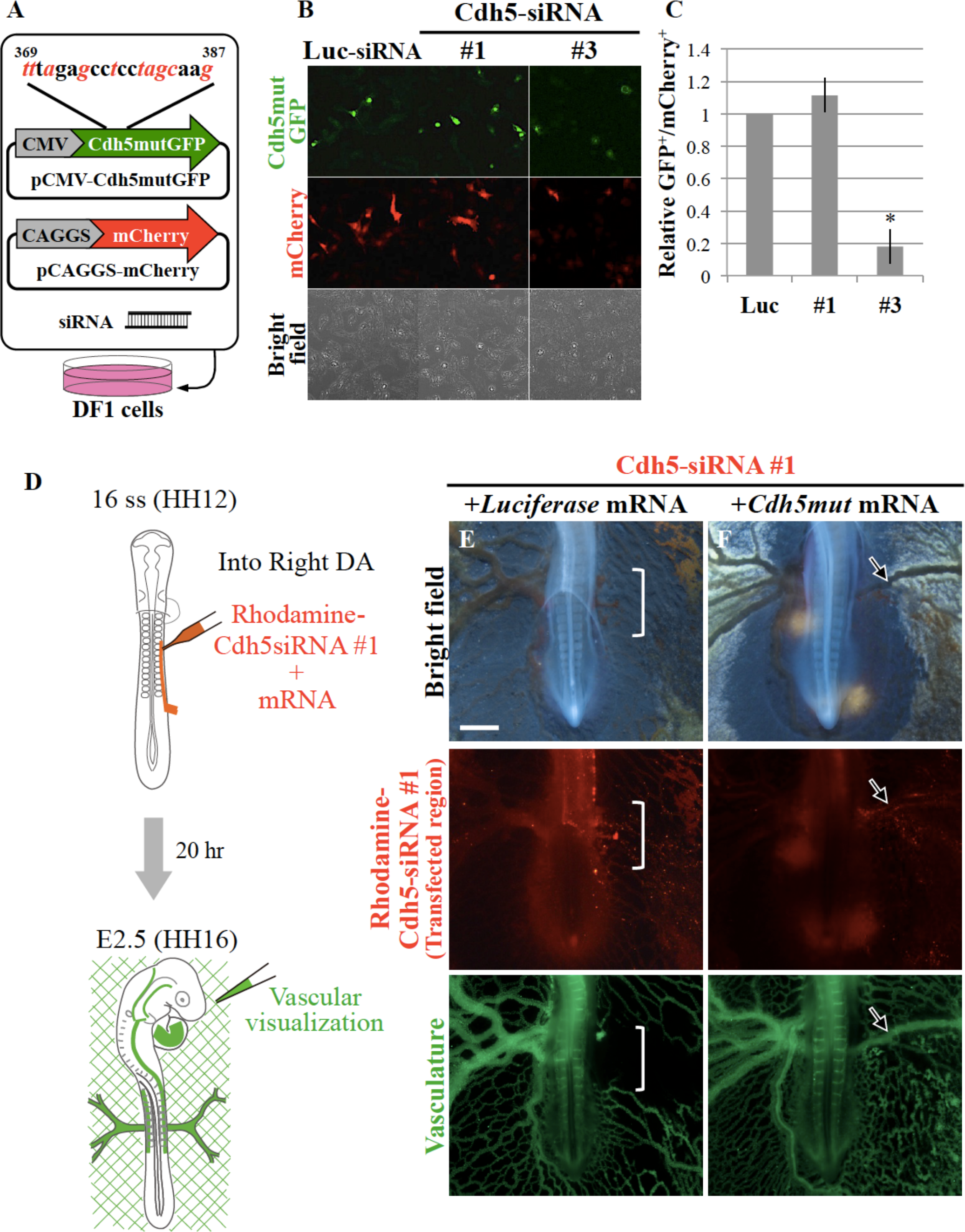
Rescue of Cdh5 siRNA-knockdown phenotypes by co-transfection with Cdh5 mRNA which was resistant to Cdh5-siRNA#1 (red letters are replaced nucleotides). (A) pCMV-Cdh5mutGFP and pCAGGS-mCherry plasmids were co-transfected into DF1 cells with each siRNA (Luc-siRNA, Cdh5-siRNA#1, and −#3), and assessed 24 hrs after transfection. (B) Cdh5mutGFP and mCherry expression in DF1 cells co-transfected with siRNAs shown on the top. Compared to the number of GFP-positive cell co-transfected with Luc-siRNA, that of Cdh5-siRNA#3 co-transfected cells was smaller. However, the number of GFP-positive cells co-transfected with Cdh5-siRNA#1 was comparable to that of Luc-siRNA-cells. (C) Quantitative representation of a relative ratio of the number of GFP^+^ cells to that of mCherry^+^ cells (n=4 each). Thus, Cdh5-siRNA#1 failed to knockdown Cdh5mutGFP, whereas Cdh5mutGFP was susceptible to Cdh5-siRNA#3. Error bars represent SD (standard deviation). \**P* < 0.001. (D) A cocktail containing rhodamine-labeled Cdh5-siRNA#1, mRNAs of *Luciferase* or *Cdh5mut*, and Lipofectamine 2000 was injected into the R-DA of 16 ss embryos, which were assessed 20 hrs transfection (E2.5) with vascular visualization by ink perfusion (green). (E, F) Phenotypes of embryos co-transfected with Cdh5-siRNA#1 and mRNA shown on the top. White brackets indicate transfected area (rhodamine-positive). (E) Embryos co-transfected with Cdh5-siRNA#1 and *Luciferase* mRNA exhibited the local disruption of R-VA in the rhodamine-labeled region (n = 7/7) as seen in Fig. 4C. (F) However, such defects of R-VA by Cdh5-siRNA#1 was rescued when co-transfected with *Cdh5mut* mRNA (n = 7/10).

mRNA of the *Cdh5 mutant (Cdh5mut)*, in which the Cdh5-siRNA#1-recognizing region of the gene was mutated (Fig. 6A), was expected to be resistant to Cdh5-siRNA#1-mediated degradation, but sensitive to Cdh5-siRNA#3. To validate this resistance, DF1 cells were co-transfected with Cdh5mutGFP-encoding cDNA along with either Cdh5-siRNA#1 or Cdh5-siRNA#3 (Fig. 6A). Whereas knockdown efficiency (GFP^+^ cells/ mCherry^+^ cells) by Cdh5-siRNA#3 was similar to in Fig. 3 (0.18 ± 0.11; n=4), Cdh5-siRNA#1 failed to inhibit Cdh5mutGFP (1.11 ± 0.10; n=4) (Fig. 6B, C). Thus, Cdh5mutGFP was resistant against Cdh5-siRNA #1-mediated degradation.

Finally, we asked if *Cdh5mut* mRNA would rescue the *in vivo* vascular phenotype elicited by Cdh5-siRNA#1. Right dorsal aorta (R-DA) of 16 ss embryo was injected with a 0.8 μl cocktail containing Cdh5-siRNA#1 and *Cdh5mut* mRNA (12.5% of Lipofectamine 2000, 4 μM of rhodamine-labeled Cdh5-siRNA#1, and 360 ng/μl of *Luciferase* mRNA or 600 ng/μl of *Cdh5mut* mRNA), and vascular formation was assessed after 20 hrs at HH16 (Fig. 6D). When *Luciferase* mRNA and Cdh5-siRNA#1 were co-injected as a control, the vasculature was disrupted in the rhodamine-labeled area as seen in Fig. 4C (Fig. 6E, n=7/7). In contrast, when *Cdh5mut* mRNA was co-transfected with Cdh5-siRNA#1, overt structures of vitelline artery and yolk sac vasculature with blood circulation were recognized in the rhodamine-labeled area (Fig. 6F, n=7/10). Thus, we conclude that the vascular defects seen in Cdh5-siRNA#1-transfected embryos are caused by Cdh5-specific knockdown and not by off-target effects. The results also show that the transfection method described in this study is applicable not only for DNA/plasmid, but also for mRNA products.

## Discussion

We have demonstrated that blood flow-mediated gene transfer into developing vasculature of chicken embryos enables efficient transgenesis in a spatio-temporally controlled manner. The gene transfer is achieved by infusion with a cocktail containing DNA/RNA and Lipofectamine, the latter being a commercially available easy-to-use reagent. By exploiting stage-dependent patterns of blood flow combined with the siRNA technique, we have shown, for the first time, that endogenous gene expression of *Cdh5* can be knocked down in a site-specific manner in developing vasculature.

### Lipofectamine is useful for infusion-mediated gene transfer

We have found that among commercially available reagents commonly used for gene transfection into cultured cells, Lipofectamine 2000 and −3000, the liposomal reagents, give higher efficiency than the non-liposomal reagents, ViaFect, FuGENE HD, and SuperFect (Fig. 1, Table 1). This suggests that liposomal components are critical for the infusion-mediated transfection at least into early vasculatures *in ovo*, and this is consistent with the previous reports in which elaborately prepared liposomal constructs were used for DNA transfection into vasculatures in chicken embryos (Bollerot et al., 2006; Decastro et al., 2006). Bollerot et al. developed their original reagent by mixing lipid RPR209120 (synthesized in their laboratory) and the neutral helper lipid DOPE. Decastro et al. prepared their original reagent by mixing three lipids, cationic lipid DOSPA, DOPE, and mPEG-SS-DOPE (synthesized on their own). Both groups succeeded in transfection with infused genes into developing vasculature, and Bollerot et al. further conducted siRNA-knockdown of the co-infused exogenous *GFP* gene. These pioneering techniques, however, required high expertise to prepare reagents. In the meantime, commercially available reagents, such as Lipofectamine 2000 or −3000, have been extended to *in ovo* transgenesis of primordial germ cells (PGCs) in chickens, for which Lipofectamine-DNA cocktail is infused into blood stream (PGCs in avians are translocated by blood circulation) (Tyack et al., 2013). Indeed, we have found that Lipofectamine reagents exhibit high transfection efficiency in developing vasculature. Although we have not directly compared, the transfection efficiency by Lipofectamine used in the current study appears to be at least comparable with, if not superior to, the efficiency described in the aforementioned previous studies using liposomes prepared by their own (Bollerot et al., 2006; Decastro et al., 2006).

In our hands, whereas the transfection efficiency is comparable between Lipofectamine 2000 and Lipofectamine 3000, the former costs less than the latter. For both maximizing the efficiency of gene transfer and minimizing embryonic lethality, the concentration of Lipofectamine in the DNA-lipid cocktail is critical, which should be in the range of 12.5 % to 25 %. This concentration-sensitivity of Lipofectamine might be a reason why this popular reagent has been overlooked for *in vivo/in ovo* transfection.

### Blood flow-mediated site-specific gene manipulation in early vasculature

We were able to control targeted sites of vascular transgenesis in two different ways. One is by controlling infusion sites, which results in different areas of transfection. The other one is by controlling time points of infusion: since the early patterns of blood flow and vasculature changes drastically as development proceeds, an infused cocktail also spreads differently in a stage-dependent manner even when the cocktail is infused into the same site.

#### Differential sites of infusion

When we inject a *GFP*-Lipofectamine cocktail into the heart after the circulation is established, vasculatures in both right and left halves are equally transfected in addition to the heart (Fig. 2F). In contrast, when the cocktail is infused into the right dorsal aorta, the vasculature in only the right half of embryo/extraembryo is positive for GFP (Fig. 2I).

#### Differential time points of infusion

Our infusion-mediated gene transfer in vasculature takes advantage of the blood flow. When a cocktail is infused into the heart at 13 ss when the blood flow is about to start, the transfected area is limited (Fig. 2E). In contrast, when the cocktail is infused into the heart at 19 ss when the blood flow is more prominent than 13 ss, the infused reagents are more widely spread in the vasculature, resulting in a wide area of transgenesis (Fig. 2G).

### Vascular subtype-specific knockdown of endogenous genes using siRNAs

Last, we described that the endogenous *VE-cadherin/Cdh5* gene in vasculature can be locally knocked down by combining Lipofectamine-mediated transfection with siRNAs. The importance of Cdh5 for the vascular assembly has been appreciated since an infusion of function-blocking Cdh5 antibody causes disassembly of vessels in mice (Crosby et al., 2005). However, investigations at high resolution of the role of Cdh5 in the peripheral vasculature have largely been hampered, since transgenic animals such as mice or zebrafish, in which the Cdh5 gene is knocked out/down, exhibit early lethality, mainly because of a heart failure (Carmeliet et al., 1999; Gory-Faure et al., 1999; Montero-Balaguer et al., 2009). In the current study, we have succeeded, for the first time, in knockdown of the endogenous *Cdh5* specifically in the right half vasculature, which results in severe malformation of blood vessels without affecting the heart formation (Fig. 4C). The rescue experiments using siRNA-resistant mutant *Cdh5* mRNA indicate that the vascular deficiency elicited by Cdh5-siRNA is not by off-target means (Fig. 6F).

In summary, the method described in this study enables efficient gene transfer into early chicken vasculature without affecting heartbeat. Infusion of a cocktail with Lipofectamine, an easy-use reagent, delivers plasmid DNAs, mRNAs, and siRNAs to developing vasculatures. Vascular type-specific transfection can be achieved by adjusting infusion sites and developmental stages of embryos. This method could also be applicable to other experimental animals, facilitating investigations of the formation of peripheral vascular systems, and their roles in tissue and organ formation.

## Acknowledgments

We thank Drs. S. F. Gilbert (Swarthmore, USA) and E. Ohata (NAIST, Japan) for carefully reading the MS and a help with DNA cloning, respectively. This work was supported by JSPS KAKENHI: Grant-in-Aid for Scientific Research (B).

